# The FANCC-FANCE-FANCF complex is evolutionarily conserved and regulates meiotic recombination

**DOI:** 10.1101/2022.07.05.498687

**Authors:** Dipesh Kumar Singh, Rigel Salinas Gamboa, Avinash Kumar Singh, Birgit Walkemeir, Jelle Van Leene, Geert De Jaeger, Imran Siddiqi, Raphael Guerois, Wayne Crismani, Raphael Mercier

**Author notes:** Corresponding author: Raphael Mercier.

## Abstract

At meiosis, programmed meiotic DNA double-strand breaks are repaired *via* homologous recombination, resulting in crossovers (COs). From a large excess of DNA double-strand breaks that are formed, only a small proportion gets converted into COs because of active mechanisms that restrict CO formation. The Fanconi anemia (FA) complex proteins AtFANCM, MHF1, and MHF2 were previously identified in a genetic screen as anti-CO factors that function during meiosis in *Arabidopsis thaliana*. Here, pursuing the same screen, we identify FANCC as a new anti-CO gene. FANCC was previously only identified in mammals because of low primary sequence conservation. We show that FANCC, and its physical interaction with FANCE-FANCF, is conserved from vertebrates to plants. Further, we show that FANCC, together with its subcomplex partners FANCE and FANCF, regulates meiotic recombination. Mutations of any of these three genes partially rescues CO-defective mutants, which is particularly marked in female meiosis. Functional loss of FANCC, FANCE, or FANCF results in synthetic meiotic catastrophe with the pro-CO factor MUS81. This work reveals that FANCC is conserved outside mammals and has an anti-CO role during meiosis together with FANCE and FANCF.

## Introduction

Large-scale exchange of genetic material between homologous chromosomes in the form of meiotic crossovers (COs) generates new allelic combinations in the sexual progeny of eukaryotes. COs are also required for the correct segregation of chromosomes at the first meiotic division in most species. This is likely why a mechanism exists to ensure an “obligate” crossover per chromosome pair, per meiosis. Meiotic recombination is initiated by the formation of a large number of programmed DNA double-stranded breaks (DSBs), a minority of which are repaired as COs. Two pathways contribute to CO formation, defining two classes of COs. Class I COs depend on a group of proteins called ZMMs, an acronym derived from seven proteins initially described in *Saccharomyces cerevisiae* (Zip1-Zip4, Msh4-5, Mer3) (Pyatnitskaya, Borde, and De Muyt 2019; Borner, Kleckner, and Hunter 2004), and account for most of the COs. Class II COs, which account for a minority of COs in most eukaryotes including mammals and plants, involve notably the MUS81 nuclease (Youds and Boulton 2011; de los Santos et al. 2003).

In *Arabidopsis thaliana*, a mutation in any member of the *ZMMs* causes a drastic decrease in CO number, and notably the loss of the obligate crossover, with only a few residual COs formed by the class II pathway, leading to chromosome mis-segregation and quasi-sterility. A forward genetic screen for restoration of fertility of *zmm* mutants identified a series of genes that actively limit CO formation in Arabidopsis. These factors specifically limit class II COs and act through three mechanisms. The first anti-CO pathway involves proteins from the Fanconi anaemia (FA) pathway, FANCM (Crismani et al. 2012), MHF1, and MHF2 (Girard et al. 2014). These three proteins have been shown to physically interact in humans, along with FAAP24, which has not been identified in Arabidopsis (Singh et al. 2010; Tao et al. 2012). The second anti-CO mechanism involves RECQ4, RMI1, and TOP3a (Seguela-Arnaud et al. 2017), and the third, the proteins FIGL1 and FLIP (Girard et al. 2015; Fernandes, Duhamel, et al. 2018; Kumar et al. 2019). These three mechanisms contribute in parallel to limiting class II COs (Fernandes, Seguela-Arnaud, et al. 2018).

The FA pathway is comprised of at least 23 protein subunits in human cells and some, but not all, of them are widely conserved in eukaryotes, including plants. It is traditionally known for its role in inter-strand crosslink repair in somatic cells in humans, with emerging roles in replication fork protection (Kolinjivadi, Crismani, and Ngeow 2020). The FA pathway is heavily studied because its proper function is required to prevent serious human disease: FA functions as a tumor suppressor and mutation of FA pathway factors causes the rare condition Fanconi anemia. FA core complex proteins have been classified into three groups on the basis of their molecular roles: (i) The core complex encompasses the largest group of proteins and is installed at DNA damage sites or stalled replication forks, with FANCM acting as an anchor. MHF1-MHF2, a heterodimeric protein complex, promotes FANCM recruitment at the site of DNA damage (Yan et al. 2010; Singh et al. 2010). (ii) The FA-ID2 complex is recruited and monoubiquitinated by the core complex at the DNA damage site (Smogorzewska et al. 2007), while it antagonizes the recruitment of FA proteins to chromatin in the absence of DNA damage (Lopez-Martinez et al. 2019). (iii) FA/HR complex proteins are downstream partners that are considered to function independently of the above two groups (Kottemann and Smogorzewska 2013; Deakyne and Mazin 2011).

Structural studies (Shakeel et al. 2019; Swuec et al. 2017; Huang et al. 2014) have demonstrated that seven subunits of the core complex (FANCA, FANCB, FANCC, FANCE, FANCF, FANCG, FANCL), and two FA-associated proteins (FAAP20 and FAAP100) form three different subcomplexes: (i) FANCB-FANCL-FAAP100 (BL100), (ii) FANCC-FANCE-FANCF (CEF), and (iii) FANCA-FANCG-FAAP20 (AG20)(Huang et al. 2014; Rajendra et al. 2014). The ring finger domain of FANCL acts as an E3 ubiquitin ligase and its two associated proteins FANCB and FAAP100 are organized as a catalytic module (Huang et al. 2014; Shakeel et al. 2019). It has been proposed that FANCA and FANCG form a chromatin-targeting module, while FANCC, FANCE, and FANCF organize to establish a substrate-recognition module (Shakeel et al. 2019). FANCF acts as a bridge between FANCC and FANCE (Leveille et al. 2004; Shakeel et al. 2019). FANCM interacts with the core complex through FANCF (Deans and West 2009; Huang et al. 2014), demonstrating that the substrate-recognition module is an important component of the FA core complex.

In this study, extending a previously described forward genetic *zmm* suppressor screen (Crismani et al. 2012; Girard et al. 2014; Fernandes, Duhamel, et al. 2018; Seguela-Arnaud et al. 2017; Girard et al. 2015), augmented by complementary approaches, we demonstrate that the CEF complex is evolutionarily conserved from mammals and show that it is a novel meiotic anti-CO factor.

## Results

### Identification of a novel *zmm* suppressor

CO-deficient *zmm* mutants display a > 90% reduced seed set in Arabidopsis, which correlates with shorter fruit, due to random segregation of chromosomes in meiosis. Therefore, inactivating anti-CO genes in a *zmm* mutant leads to an increase in CO number, resulting in improved chromosome segregation, restored fertility and longer fruits. Here, we extended a forward genetic screen for *zmm* mutants exhibiting increase in fruit length following EMS mutagenesis. The screen was previously performed on five *zmm* mutants (*hei10, zip4, shoc1, msh5* and *msh4*), in a total of ∼7,000 lines and identified 59 mutants with restored fertility, among which 58 are mutated in one of the previously identified anti-CO genes (Table 1, Table S1). (Girard et al. 2014; Girard et al. 2015; Crismani et al. 2012; Fernandes, Duhamel, et al. 2018; Seguela-Arnaud et al. 2017). In this study, we focused on the last *zmm* suppressor mutation that increased the fertility of a *msh4* mutant (cshl_GT14269, L*er* genetic background). Genetic mapping delimited the causal mutation to a 0.47MB region on chromosome 3 (21452882-21919909 in the Ler assembly)(Zapata et al. 2016), and whole genome sequencing identified a candidate mutation in the fourth exon donor splicing site of the At3g60310 gene (G>A 3_21918909 in the Ler assembly, corresponding to 3_22288z888 in Col TAIR10). We show below that At3g60310 encodes the Arabidopsis FANCC ortholog. Three independent T-DNA alleles (*fancc-2 N542341*, *fancc-3 N1007952* and *fancc-4 N626745*, Col background) were able to enhance the fertility of *msh4*, from 4.5 to >24 seeds per fruit (Figure 1 and Supplementary Figure S1, S2), confirming the identification of the causal mutation in At3g60310.

**Figure 1.**
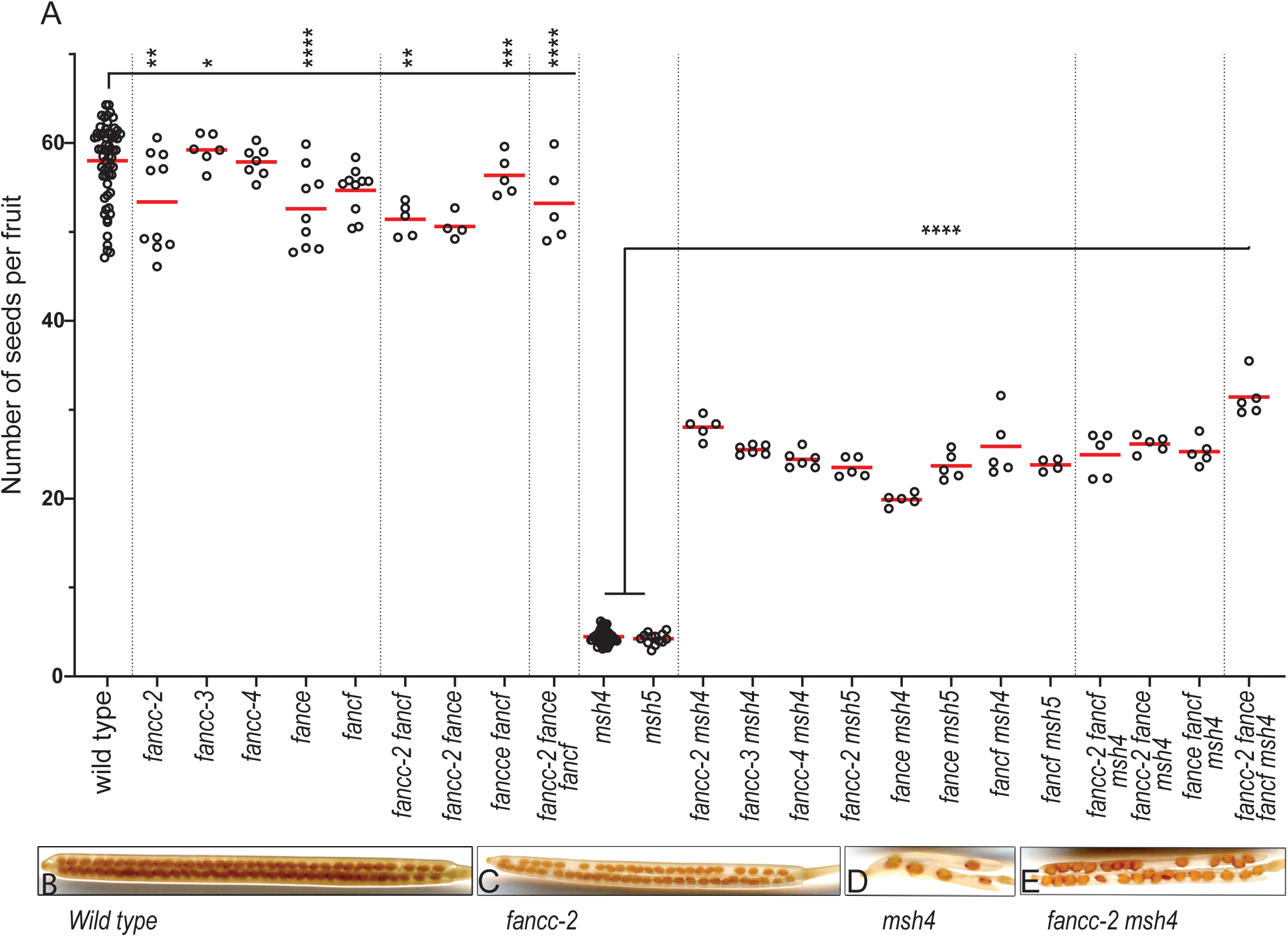
Analysis of fertility of *zmm* suppressor mutants. (A) Each dot indicates the fertility of an individual plant, measured as the number of seeds per fruit averaged on ten fruits. The mean fertility for each genotype is shown by a red bar. Each mutant was compared to sibling controls grown together, and the data of independent experiments are shown in Figure S2. Some genotypes were represented in several experiments and their data were pooled for this figure. Stars summarize the one-way ANOVA followed by Sidak test shown in Figure S2. (B–E) Representative fruits of *wild type, fancc-2, msh4*, and *fancc-2 msh4*, cleared with 70% ethanol.

**Table 1.**
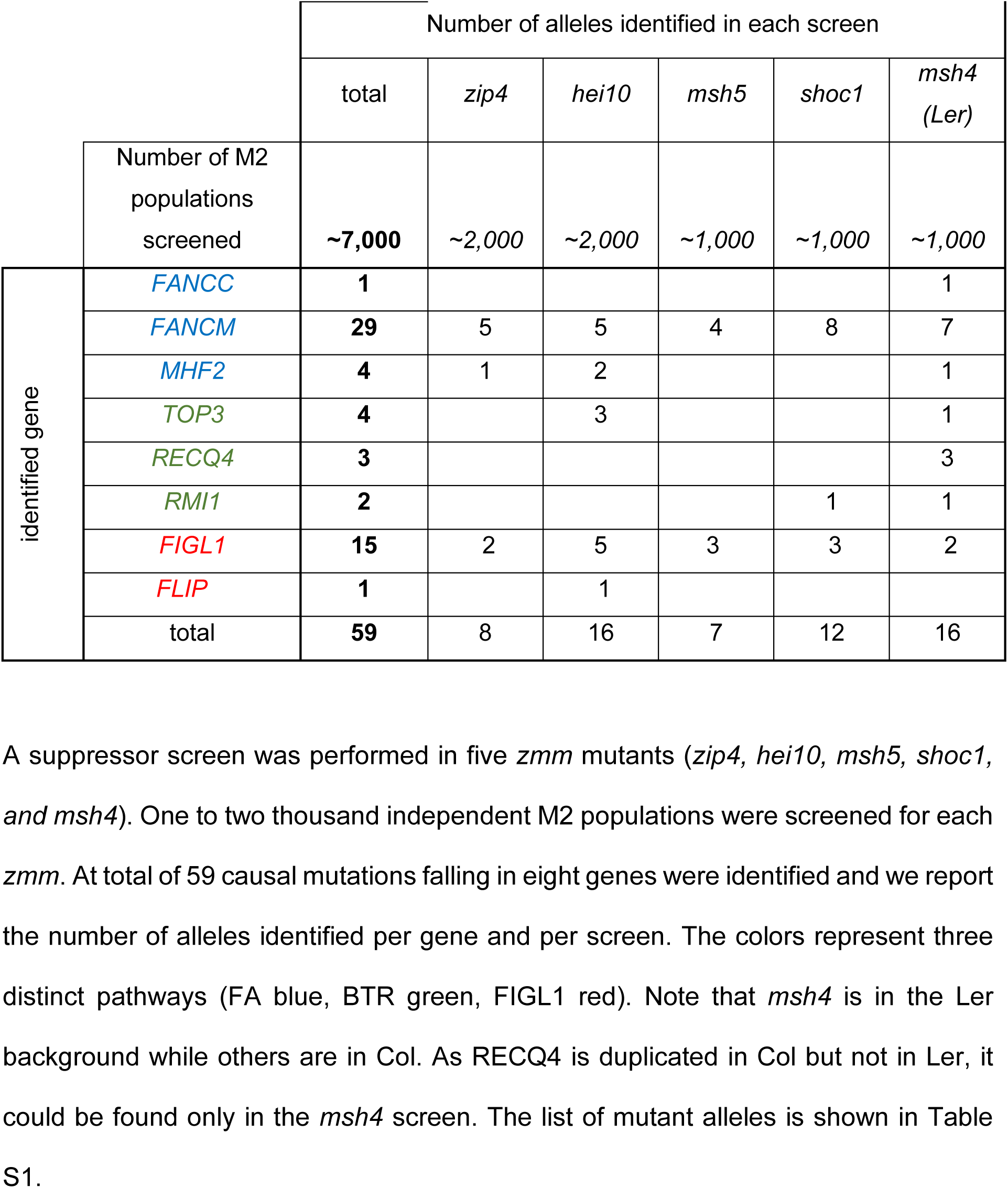
Summary of the *zmm* suppressor screen results.

|  |  | Number of alleles identified in each screen |  |  |  |  |  |
| --- | --- | --- | --- | --- | --- | --- | --- |
|  |  | total | *zip4* | *hei10* | *msh5* | *shoc1* | *msh4*<br>(Ler) |
| Number of M2 populations screened |  | ~7,000 | ~2,000 | ~2,000 | ~1,000 | ~1,000 | ~1,000 |
| identified gene | *FANCC* | 1 |  |  |  |  | 1 |
|  | *FANCM* | 29 | 5 | 5 | 4 | 8 | 7 |
|  | *MHF2* | 4 | 1 | 2 |  |  | 1 |
|  | *TOP3* | 4 |  | 3 |  |  | 1 |
|  | *RECQ4* | 3 |  |  |  |  | 3 |
|  | *RMI1* | 2 |  |  |  | 1 | 1 |
|  | *FIGL1* | 15 | 2 | 5 | 3 | 3 | 2 |
|  | *FLIP* | 1 |  | 1 |  |  |  |
|  | total | 59 | 8 | 16 | 7 | 12 | 16 |

### FANCC is conserved in plants

Standard sequence similarity analysis failed to find any homology of the protein encoded by At3g60310 outside of plants, or with proteins of known function (Stanley et al. 2016). Using the HHpred remote homology detection server (Zimmermann et al. 2018). (Soding 2005), it was possible to identify a potential match with human FANCC (XP_011516668) despite both proteins sharing only 16% primary sequence identity (HHpred probability of 94%). To test the hypothesis that At3g60310 is an ortholog of FANCC, we analyzed the physical contacts between human FANCC and the human FANC complex, the cryo-EM structure of which was recently determined at 3.1 Å (Wang et al. 2021). Figure 2A illustrates that human FANCC (pale green subunit) is in direct physical contact with three subunits of human FA core complex, FANCE (light pink), FANCF (light blue) together with the ubiquitin E3 ligase FANCL (yellow). Given that FANCC, FANCE, and FANCF are known to constitute a stable sub-complex in humans, we tested the possibility that At3g60310 was a dedicated partner of *A. thaliana* FANCE (Girard et al. 2014) (Q9SU89_ARATH) and FANCF (F4K7F0_ARATH) using the Alphafold2 prediction method (Jumper et al. 2021). Alphafold2 was recently shown to perform well when predicting structures of proteins and whether two proteins interact with each other (Evans et al. 2021). Using the AlphaFold2 method trained on multimers (Mirdita, Ovchinnikov, and Steinegger 2021), we obtained a model of the complex with the three *A. thaliana* subunits with reliability scores above the confidence threshold of 50 and 0.5 for pLDDT and ptmscore, respectively (pLDDT of 72.6 and ptmscore of 0.67) (Supp. Figure S3). Interestingly, At3g60310 was predicted to form a complex with AtFANCE and AtFANCF with a similar arrangement to that observed for the corresponding orthologs in the human FANC complex (Figure 2B). As a support for the reliability of the model, the surface patches of At3g60310 involved in the interaction with both subunits were among the most conserved regions (Figure 2C, Supp Figure S4,S5). The N-terminal domain of FANCE is found well anchored in the central region of At3g60310/FANCC with low predicted error for the accuracy of the interface modelling (Supp Figure S3). In contrast, the C-terminal domain does not exhibit a strong co-evolutionary signal in the region where it binds to At3g60310/FANCC. The five models generated for FANCE do not converge on a single conformation for the C-terminal domain, although due to the tethering of the N-terminal domain, this domain tends to cluster in the same region as the surface where the FANCL E3 Ub ligase binds to FANCC. One hypothesis could be that upon binding of the E3 ligase FANCL, the C-terminal domain of FANCE is displaced from the position shown in Figure 2B to allow FANCL binding, which is potentially coupled with the recruitment of FANCD2 substrate by the FANCE C-terminal domain (Polito et al. 2014).

**Figure 2.**
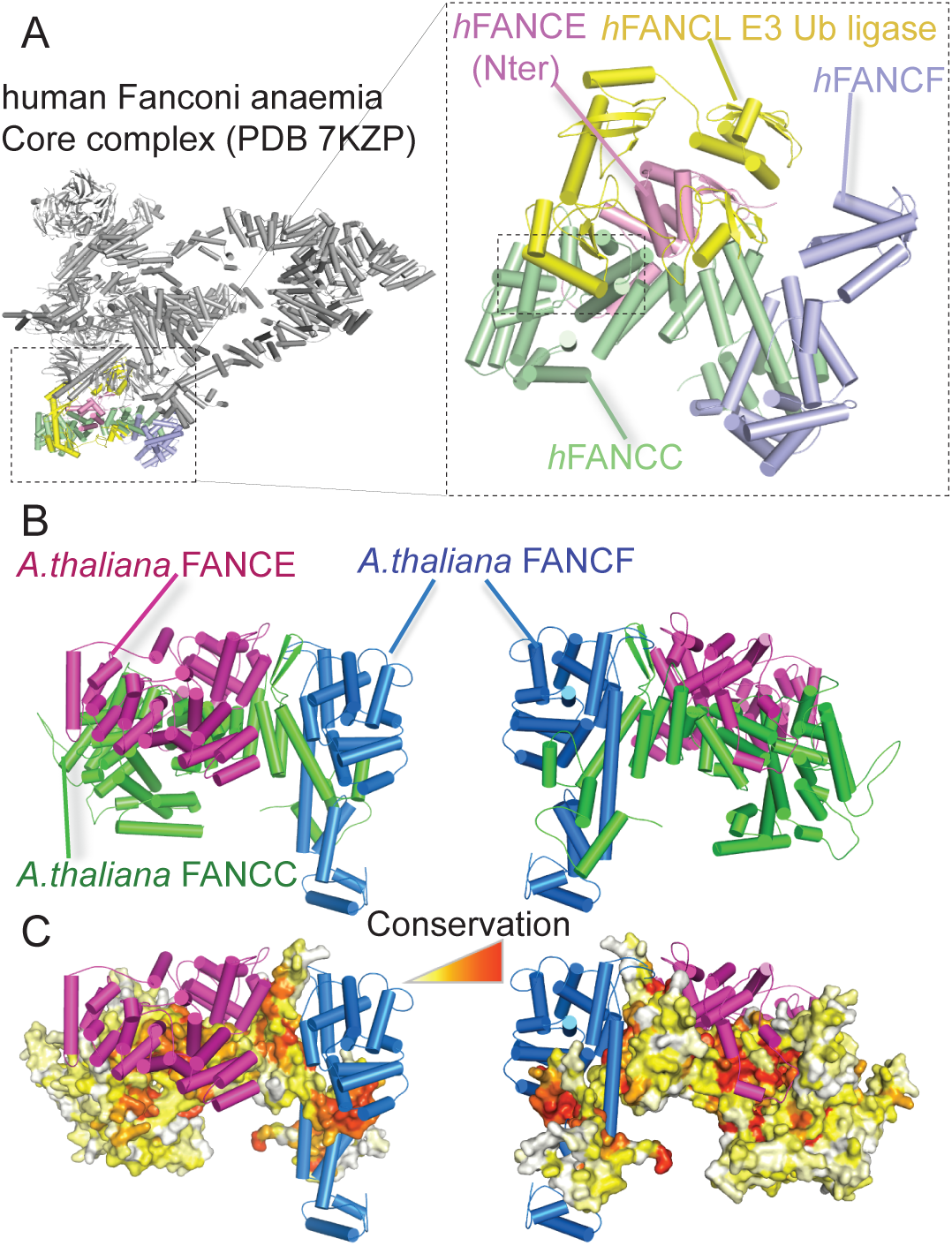
Structural analysis of the experimental human and modeled *A*. *thaliana* FANCC-FANCE-FANCF complexes. (A) Structural representation of the human FANC core complex (PDB:7KZP) (Wang et al. 2021) Most of the subunits are shown in gray with the exception of those in direct contact with the human FANCC (light green), namely hFANCE (light pink), hFANCL (yellow) and hFANCF (light blue). A zoomed-in view of the four subunits is shown in the inset on the right with the contact region between hFANCL and hFANCC highlighted by a dotted rectangle. (B) AlphaFold2 structural model of the AtFANCC-AtFANCE-AtFANCF complex represented as a cartoon in two orientations with a dotted square indicating the C-terminal domain of FANCE located in a region of FANCC that directly binds to the FANCL subunit in the human FANC core complex structure. (C) Same view as (B) with AtFANCC shown as a surface and colored according to conservation from white to red for the least to most conserved positions. Pymol software was used to draw the different structures (The PyMOL Molecular Graphics System, Version 2.0 Schrödinger, LLC).

Next, we performed an unbiased search for interacting partners of At3g60310 using pull-down protein purification coupled with mass spectrometry. We used overexpressed GSrhino-tagged At3g60310 as a bait in Arabidopsis suspension cell culture (Van Leene et al. 2019). After filtering copurified proteins for false positives, we recovered peptides from At3g60310 itself and a series of additional proteins in three replicate experiments (Table 2). Strikingly, all four co-purified identified proteins were Arabidopsis homologs of members of the FA complex, FANCE, FANCL, FANCM, and MHF2 (Table 2). Further, a yeast two-hybrid assay confirmed direct interactions of At3g60310 with FANCE and MHF2. FANCE, FANCF and MHF2 also interacted with each other in yeast two-hybrid (Figure S6). Altogether, this demonstrates that At3g60310 encodes the FANCC protein in Arabidopsis, which we term AtFANCC.

**Table 2.** Pull-down protein purification using At3G60310/FANCC as bait.

| Gene Id | Protein name | CGSrhino PD1 | CGSrhino PD2 | CGSrhino PD3 |
| --- | --- | --- | --- | --- |
| At3G60310 | FANCC | 19 | 18 | 15 |
| AT4G29560 | FANCE | 6 | 4 | 5 |
| AT5G65740 | FANCL | 5 | 4 | 4 |
| At1G78790 | MHF2 | 2 | - | 3 |
| AT1G35530 | FANCM* | 2 | - | - |

Three replicates of pull-down purifications (IP1, IP2 and IP3) followed by mass
spectrometry were performed using FANCC as a bait. After filtering (see material and
methods), the number of specific peptides is reported for each identified protein. \*Only
identified in one experiment. MS data are shown in table S4.

### FANCC is conserved in distant eukaryotic lineages

Using PSI-BLAST searches, AtFANCC orthologs could be detected in most plants (Figures S4 and S5). In-depth analyses using either PSI-BLAST or HHpred failed to detect any homolog in more distantly related green algae such as *Chlamydomonas*, although a FANCE homolog can be detected in *Chlamydomonas reinhardtii*. In metazoans, a previous bioinformatics analysis performed on model species for all the genes of the FANC core complex, noted that several species were missing a FANCC homolog although having a FANCE ortholog (Stanley et al. 2016). We revisited this study using the most recent sequence databases and PSI-BLAST searches starting from human FANCC. Interestingly, five iterations of PSI-BLAST were required to retrieve the first plant ortholog (in the monocot *Spirodela intermedia*), which enabled the retrieval of all the same plant orthologs identified from AtFANCC. After 15 iterations, the search nearly converged with about 2,100 FANCC homologs, highlighting the existence of FANCCs in early branching metazoans such as *Nematostella vectensis* (XP_032241565) and *Ciona intestinalis* (XP_002129616) that were not found previously. In insects, orthologs could also be detected in Hymenoptera (ants and bees) but neither in Diptera (Drosophila) nor in Lepidoptera (Bombyx). Consistently, repeating the PSI-BLAST search with human FANCE or FANCF as queries, a similar distribution of homologs was found in insects. Homologs of FANCE and FANCC could be detected in specific fungal lineages such as *Rhizopus azygosporus* (corresponding to hypothetical proteins RCH90546.1 and RCH79564.1, respectively). A reciprocal HHpred analysis comparing these genes against the human database confirmed they were remote homologs of FANCE and FANCC (HHpred probability score of 100% and sequence identities of 21% and 15%, respectively), suggesting that certain fungal lineages did not lose these FANC complex subunits.

### *Atfancc, Atfance,* and *Atfancf* mutations increase fertility and bivalent formation of crossover-deficient *zmm* mutants

As the mammalian FANCC was shown to form a structural and functional module with FANCE and FANCF ((Shakeel et al. 2019; Swuec et al. 2017; Wang et al. 2021), we explored their potential meiotic roles in parallel. A FANCE homolog was previously described in Arabidopsis (Girard et al. 2014), and the AT5G44010 gene was annotated as *AtFANCF* in Araport11 because of sequence similarity with the mammalian *FANCF* (Cheng et al. 2017). Single mutants in each of these genes did not show growth or developmental defects but had a slight decrease in fertility (Figure 1, Figure S2). Meiotic chromosome spreads in *Atfance* and *Atfancf* single mutants revealed the presence of univalents at low frequencies, showing that some chromosome pairs lack COs (Figure S7, Figure 3). Combination of *fancc, fance* and *fancf* mutations did not reveal any developmental defects or enhanced sterility and meiotic defects (Figures 1,3). This suggests that, consistent with the pull-down and Y2H data, FANCC, FANCE, and FANCF act together at meiosis, playing a role in ensuring the obligate CO.

**Figure 3.**
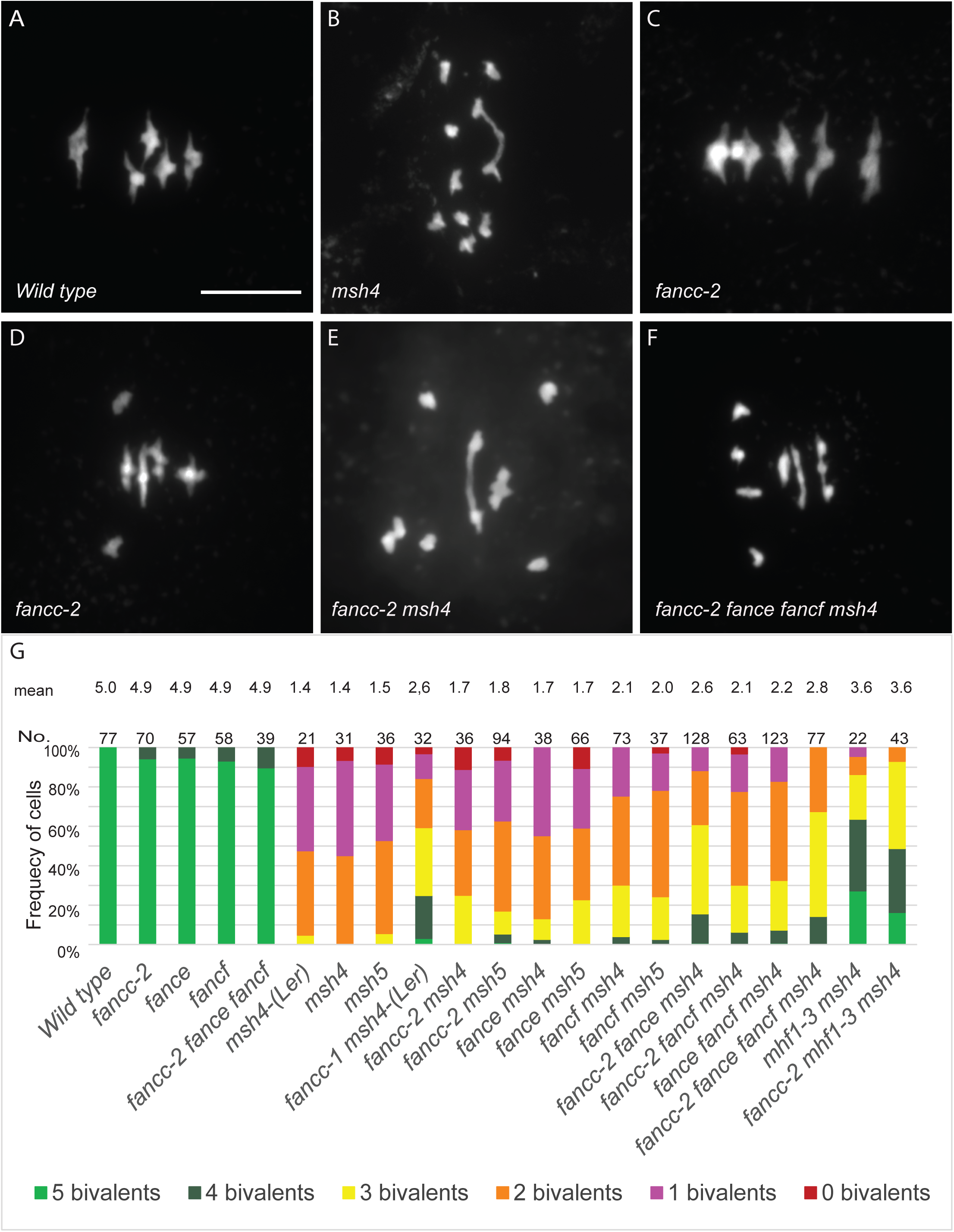
Metaphase chromosome spreads of male meiocytes. (A) Wild type with five bivalents, (*B*) *msh4* with one bivalent, (C) *fancc-2* with four bivalents and one pair of univalents. (D) *fancc-2* with five bivalents (*E*) *fancc-2 msh4* with two bivalents (F) *fancc-2 fance fancf msh4* with two bivalents. Scale bar, 10 µm. (G) Quantification of bivalents at metaphase I. The proportion of cells with 0–5 bivalents is shown with a color code. The number of analyzed cells and the average bivalent number per cell is shown for each genotype. All the genotypes are in the Col background, except when Ler is mentioned.

Mutation of *FANCC*, or *FANCE*, or *FANCF* significantly restored the fertility of the *zmm* mutants *msh4* and *msh5*, increasing the seed set more than fourfold (Sidak test, p<10^-6^) (Figure1, Figure S2). In the Ler background, chromosome spreads of male meiosis showed an increase of bivalent frequency in *fancc-1 msh4* compared to *msh4* (p<0.001) (Figure 3). In comparable experiments in the Col background, *fancc*, *fance*, or *fancf* individual mutations barely increased bivalents in *msh4* (p=0.12, 0.12 and 0.0002, respectively) (Figure 3). Combining Col *fance, fance,* and *fancf* mutations in *msh4* further restored bivalent formation to reach an average of 2.8 bivalents/cell compared to 1.4 in *msh4* (p<0.0001) and led to a slightly higher fertility increase compared to *fancc msh4* (p=0.04). Altogether, this suggests that FANCC, FANCE, and FANCF limit CO formation in a partially redundant manner. Note that this restoration of bivalent formation is lesser than that obtained through mutation of *FANCM* (5 bivalents; (Crismani et al. 2012)), MHF1 or MHF2 (3.6; Figure 3G, (Girard et al. 2014)), suggesting that the FANCC-FANCE-FANCF module has a supporting role in limiting a portion of the COs prevented by FANCM-MHF1-MHF2. The *fancc* mutation did not further restore bivalent formation in *mhf1 msh4* (t-test p=0.56), suggesting that FANCC acts in the same anti-CO pathway as MHF1 (Figure 3G).

### FANCC, FANCE, and FANCF regulate meiotic crossover formation

Intriguingly, in the above experiments, we found that *fancc*, *fance,* and *fancf* increased the fertility of *zmm* mutants (*msh4 and msh5*), but that the increase of bivalent number in male meiotic cells was less robust than the seed set suggested. As fertility in the self-pollinating plant Arabidopsis depends on both male and female meiosis, this may suggest that the role of FANCC, FANCE, and FANCF in limiting COs is more critical in female meiosis than in male meiosis. To check this, we used a test line for recombination (FTL420), which contains two transgenes conferring expression of GFP and RFP in the seed coat and defining a 5-Mb interval of the sub-telomeric region on chromosome 3 (Ziolkowski et al. 2015; Melamed-Bessudo et al. 2005). Crossover frequency was measured for males and females separately, through reciprocal crosses with wild-type plants, and in selfing (figure 4, table S 2). In females, recombination was significantly increased (Fisher test, p<0.0001) in *fancc* and *fancc fance fancf* compared to wild type, confirming the anti-CO function of *FANCC.* In males, crossover frequency was not increased, but slightly reduced (p=0.17 for fancc and 0.015 for *fancc fance fancf*). In selfing, which combines male and female meiosis products, recombination was modestly increased in *fancc* compared to wild type (p=0.003). A similar recombination picture was observed in *fance*, *fancf*, and the triple mutant *fancc fance fancf*, suggesting that the three proteins act together in limiting meiotic crossovers.

**Figure 4.**
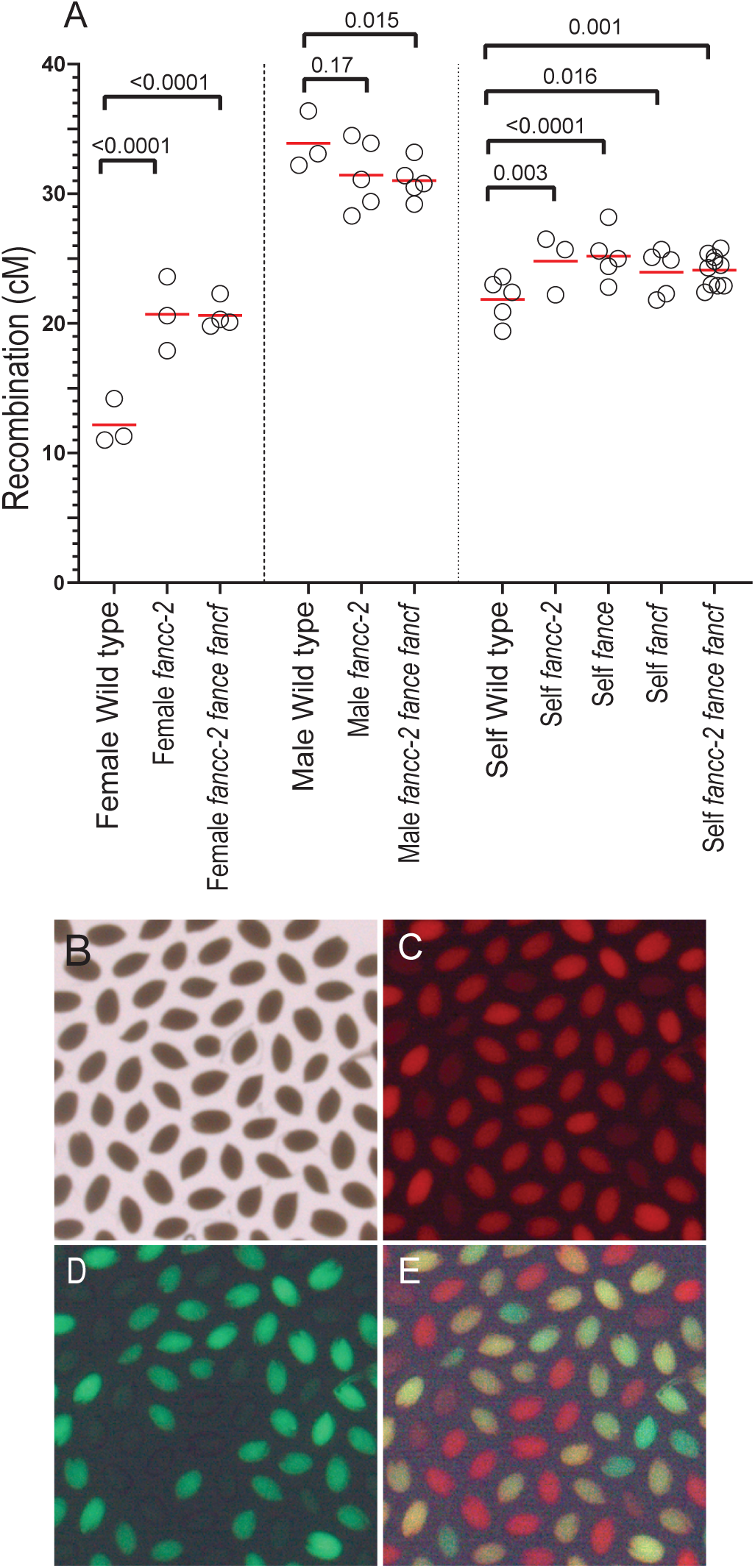
Recombination in *fancc*, *fance* and *fancf* mutant. (A) Recombination was measured in seeds produced by crosses with wild type (female and male) or after selfing. Each dot represents the recombination frequency measured in an individual plant, and the red lines show the mean. *P* values are from two-sided Fisher’s exact test on the proportion of recombined seeds. Raw data are shown in Table S2 (B–D). Representative image of seeds from a 420/++ hemizygote imaged under bright-field, red fluorescence channel, green fluorescence channel, and merged fluorescence.

### *Atfancc*, *Atfance*, and *Atfancf* exhibit chromosome fragmentation in the *mus81* background

Because of the roles of FANCM, MHF1, and MHF2 in preventing class II COs, combining mutation in these genes with mutation of *MUS81* that catalyses class II COs, leads to chromosome fragmentation at meiosis, resulting in sterility. In addition, the *fancm mus81* double mutant shows a strong developmental defect, demonstrating the role of these two genes in somatic DNA repair (Girard et al. 2014; Crismani et al. 2012) When we combined *fancc*, *fance*, or *fancf* with the *mus81* mutation, we did not observe developmental defects. However, in double mutants with *mus81* and either *fanc-c*, *-e*, or *-f* we observed a strong reduction in fertility, measured by seed per fruit, compared to the respective single mutants (Figure 5A, S8). Meiotic chromosome spreads revealed the presence of chromosome fragments at anaphase I and subsequent stages in ∼40% of the cells of the double mutants (Figures 5 B–J, S9 A–D and F–J). This demonstrates that FANCC, FANCE, and FANCF are important for efficient DSB repair in a *mus81* background and suggests that they regulate class II CO formation but with a less critical role than FANCM and MHF1/2. The removal of all four genes – *mus81 fancc fance fancf –* did not drastically enhanced fertility defects or chromosome fragmentation compared to the double *mus81 fanc* combinations. These results support the hypothesis that all three genes, *FANCC, FANCE*, and *FANCF*, are required at meiosis to repair a subset of intermediates that can also be repaired by MUS81.

**Figure 5.**
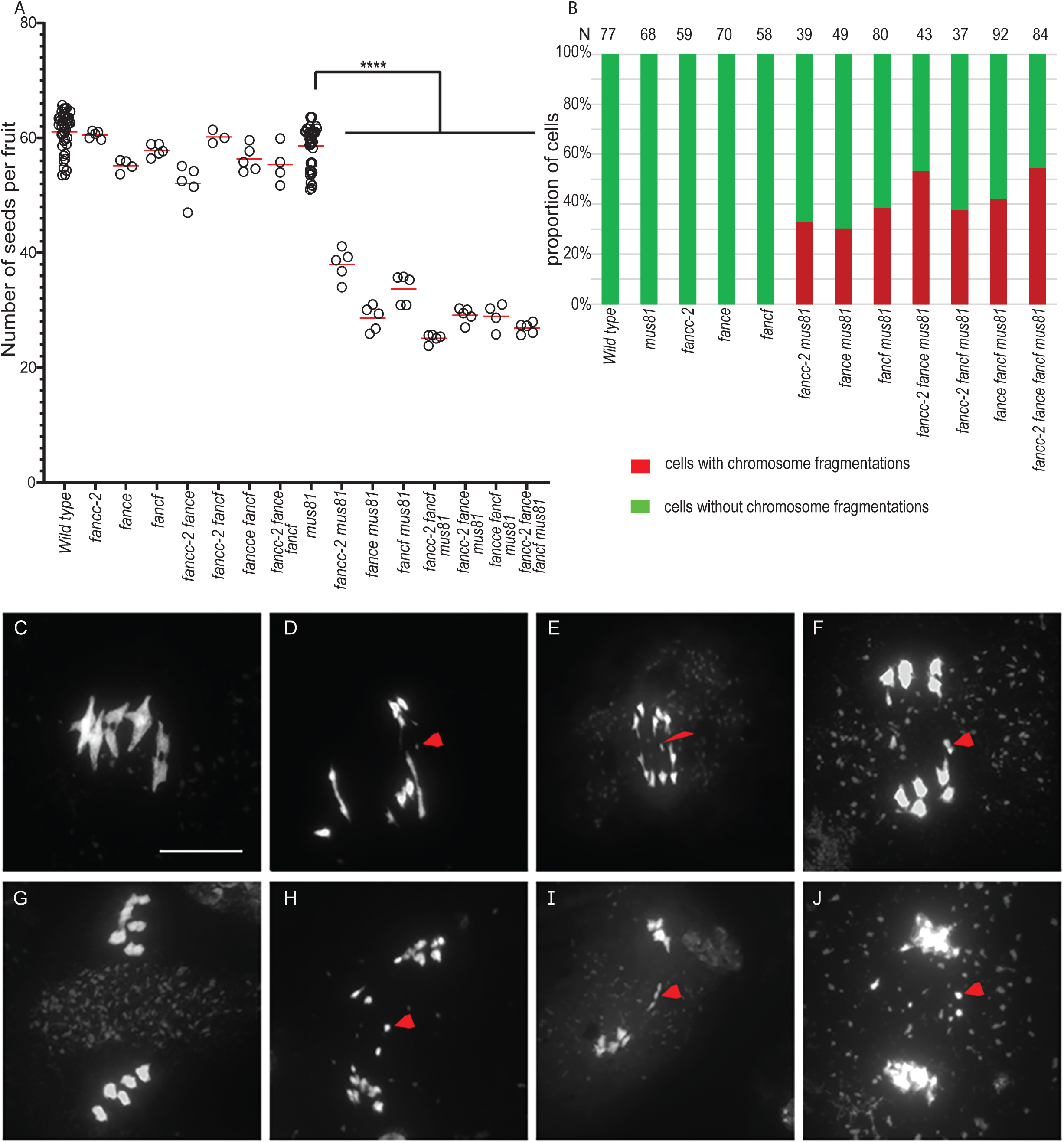
Combining *fanc and mus81* mutations leads to reduced fertility and chromosome fragmentation at meiosis. (A) Each dot indicates the fertility of an individual plant, measured as the number of seeds per fruit averaged on ten fruits. The means for each genotype are shown by red bars. Each double mutant was compared to sibling controls grown together; the independent experiments are shown in Figure S8. Wild type and *mus81* control were represented in several experiments and their data are pooled in this plot. (B) Quantification of cells with and without chromosome fragments. N=number of cells analyzed for each genotype. Stars summarize the one-way ANOVA followed by Sidak test shown in Figure S8. (C–J) Chromosome spreads of male meiocytes (Scale bar, 10 µm). Arrow heads indicate chromosome fragments. (C, G) *mus81*. (D, H) *fancc-2 mus81.* (E, I) *fance mus81* (F, J) *fancf mus81*.

## Discussion

FANCC-FANCE-FANCF constitute a stable sub-complex within the FA core complex. Based on sequence conservation, FANCE and FANCF homologs have been identified in evolutionarily distant eukaryotes such as plants (Girard et al. 2014; Cheng et al. 2017). However, despite multiple studies that systematically catalogued FA pathway protein conservation across diverse taxa, homologs of FANCC have not been identified beyond vertebrates (Zhang et al. 2009; Girard et al. 2014), suggesting that FANCC may not be conserved over large evolutionary scales. Here, combining genetics, *in vivo* pull-downs, direct protein-protein interaction studies, and structural modeling, we unambiguously identified the FANCC protein in Arabidopsis. In addition, interaction and modeling studies strongly suggest that FANCC, FANCE, and FANCF form a sub-complex in Arabidopsis as they do in vertebrates. Homologs of FANCC can also be readily identified in most other plants. As the plant and animal branches diverged very early in the eukaryotic tree of life (Burki et al. 2020), this suggests that the FA complex and notably the FANCC-E-F subcomplex was already present in the common ancestor of all living eukaryotes. The algorithm we used to detect divergent homologs succeeded in detecting the link between plant and vertebrate FANCC but failed to detect homologs in fungal lineages, except for a few species. As fungi are more closely related to animals than plants, this suggests that most of the fungal lineages have lost FANCC, or that the FANCC sequence has diverged beyond what we can recognize with current tools. Similarly, FANCC was detected in diverse animal lineages including some insects, but not in Drosophila, which can be attributed either to gene loss or to extreme divergence.

We initially identified *FANCC* because its mutation can partially restore the fertility of CO-defective *zmm* mutants, in a similar manner to previously identified anti-CO factors and notably the FA complex components FANCM, MHF1, and MHF2 (Crismani et al. 2012; Girard et al. 2014). We also found that mutation in either of the two other subunits of the FA CEF subcomplex, *fance* and *fancf*, improves the fertility of *zmm* mutants. Mutations in the three genes individually restored fertility of *zmm* to similar levels, but to a much lower level than previously obtained with *mhf1*, *mhf2* or *fancm*. Further, restoration of *zmm* fertility upon cumulative mutations in *fancc, fance,* and *fancf* remained limited. This suggests that FANCC, FANCE and FANCF together regulate meiotic recombination, but with a less critical role than FANCM, MHF1, and MHF2. We observed an increased number of bivalents at male meiosis when mutating *fancc*, *fance*, and *fancf* in *zmm* mutants, consistent with an anti-CO function. However, the increase in bivalents in males was limited compared to the observed increase in fertility, suggesting that male and female meiosis could be differently affected. We also observed a slight decrease in fertility and a low frequency of univalents in male meiocytes in single *fancc*, *fance*, or *fancf* mutants, suggesting a pro-CO function. Technical limitations prevented us from measuring the frequency of bivalents/univalents in female meiosis. However, when assessing recombination by a genetic assay in *fancc* and *fancc fance fancf*, we observed a large increase in recombination in females and a small decrease in males. Altogether, we propose that FANCC-E-F regulates meiotic recombination, with a predominant anti-CO function in females, explaining the capacity of their mutation to restore the fertility of *zmm* mutants.

Similar to FANCM and MHF1/MH2, we propose that FANCC-E-F prevents the formation of class II COs that are catalyzed by MUS81. Indeed, combining any of *fancc*, *fance* or *fancf* with the *mus81* mutation led to chromosome fragmentation at meiosis and reduced fertility (Figure 5). Combining the three mutations (*fancc fance fancf*) together had only a slightly increased effect compared to single mutants in the capacity to increase fertility and bivalents of *zmm* mutants (Figure 1 and 3) or for synthetic meiotic catastrophe and reduced fertility when combined with *mus81* (Figure 5). Further, the recombination assay did not detect differences between the single *fancc* and the triple *fancc fance fancf* (Figure 4), suggesting that FANCC-FANCE-FANCF act together in regulating recombination. In all experiments, the observed effects – even when combining the three *fancc fance fancf* mutations – are weaker than observed with MHF1/2 or FANCM (Crismani et al. 2012; Girard et al. 2014). Further, the *fancc-2* has no additive effect with *mhf1* (figure 3). Altogether, this shows that FANCC-FANCE-FANCF acts in the same anti-CO pathway as FANCM-MHF1/2, as also supported by the fact that they form a stable complex *in vivo* (Table 1). We propose that FANCC-FANCE-FANCF supports FANCM activity, which unwinds recombination intermediates and directs them to non-crossover repair. We favor the hypothesis that the meiotic crossover-limiting role of Arabidopsis FANCC-E-F is distinct from the well-described somatic role of human FANC-C-E-F where it facilitates FANCD2-FANCI mono-ubiquitination in inter-strand crosslink repair. This hypothesis is supported by the following lines of evidence: 1) There is no detectable crossover-limiting role of neither of the Arabidopsis orthologues of the catalytic component of the human FA core complex – the E3 RING ligase, FANCL – and its substrate FANCD2-FANCI (Girard et al. 2014; Kurzbauer et al. 2018); 2) Human FANCM has well-described functions distinct from the FA core complex (Walden and Deans 2014; Deans and West 2009; Ito and Nishino 2021) that are associated with remodeling branched DNA structures. The FANCC-E-F complex may act to stabilize or support the activity of FANCM in performing its function of branched molecule dissolution during meiotic DSB repair.

## Materials and methods

### Genetic material

The following Arabidopsis lines were used in this study: *fancc-2* (N542341), *fancc-3*(N1007952), *fancc-4* (N626745), *fance* (N553587) (Girard et al. 2014) *fancf* (N457070) msh4 (N636296(Higgins et al. 2008), *msh5-2* (N526553)(Higgins et al. 2008), *mus81*-2 (N607515) (Berchowitz et al. 2007), and *mhf1-3*(N576310) (Girard et al. 2014). All the T-DNA mutants were obtained from the NASC.

### Genetic analysis

The msh4 suppressor *Atfancc* was sequenced using IIlumina technology at The Genome Analysis Centre, Norwich, UK, and mutations were identified using ler 1 assembly as reported for the MutDetect pipeline (Girard et al. 2014; Schneeberger et al. 2011). The identified causal mutation in fancc-1 was a G to A substitution at position chr3: 21918909 (Ler-0 TAIR10 assembly) equivalent to position chr3: 22288888 in the Columbia (TAIR10) genome. The primers used for genotyping are listed in Table S3. Siliques were fixed in 70% ethanol for at least two days and scanned for seed counting manually on images. Fertility was assessed by counting seeds per fruit on a minimum of five plants and ten fruits from each plant.

### Sequence analyses

Sequences of *A*. *thaliana* At3g60310/AtFANCC, FANCE (Q9SU89_ARATH) and FANCF (F4K7F0_ARATH) were used as input for the HHpred remote homology detection server against different eukaryotic profile databases (Zimmermann et al. 2018)(Soding 2005) and as queries of PSI-BLAST searches (Altschul et al. 1997) against the nr database. Full-length sequences of FANCC orthologs were retrieved and re-aligned with mafft (Katoh and Standley 2013) and the multiple sequence alignment was represented using JalView (Waterhouse et al. 2009). The phylogenetic tree of the FANCC orthologs in plants was generated using the FANCC MSA as a query of the PhyML 3.0 server (Dereeper et al. 2008) with standard estimated options, an approximate likelihood-ratio test to estimate the bootstrap values (SH-like), and the Jones-Taylor-Thornton substitution model with four substitution rate categories. The calculated tree was represented using the iTOL server (Letunic and Bork 2021).

### Structural modeling

Sequences of *A*. *thaliana* At3g60310/AtFANCC, FANCE (Q9SU89_ARATH) and FANCF (F4K7F0_ARATH, the) query sequences of each subunits were used as input for the MMseqs2 homology search program (Steinegger and Soding 2017) to generate a multiple sequence alignment (MSA) against the UniRef30 clustered database for each of the FANC complex subunits (Mirdita et al. 2017). The calculated full-length sequences of the orthologs were retrieved and re-aligned with mafft (Katoh and Standley 2013). MSAs of FANCC, FANCE and FANCF were then concatenated, matching the sequences of the same species resulting in paired alignments, which were combined with the unpaired sequences for those species that could have not have been matched. The resulting paired plus unpaired concatenated MSA was used as input to generate five structural models of the FANCC-FANCE-FANCF complex using a local version of the ColabFold interface (Mirdita, Ovchinnikov, and Steinegger 2021) running three iterations of the Alphafold2 algorithm (Jumper et al. 2021) trained on the multimer dataset (Evans et al. 2021) on a local HPC equipped with NVIDIA Ampere A100 80Go GPU cards. The five models converged toward similar conformations and obtained high confidence and quality scores with pLDDTs in the range [79.1, 85] and [72.6, 80.2] and pTMscore in the range [0.64, 0.662]. The model with highest pTMscore was relaxed using rosetta relax protocols to remove steric clashes constrained by the starting structure using the -relax:constrain_relax_to_start_coords option (Leman et al. 2020), and the model with the lowest rosetta global energy was used for structural analysis. Conservation analysis mapped at the structure of the model were performed using the ConSurf server (Ashkenazy et al. 2016).

### Cytological techniques

Meiotic chromosome spreads on anthers were performed as previously described (Ross, Fransz, and Jones 1996). Chromosomes were stained with DAPI (1 μg/ml) Images were acquired and processed using a ZEISS microscope (AXIO-Imager.Z2) under a 100× oil immersion objective with ZEN software and figures were prepared using Adobe Photoshop.

### Yeast two-hybrid and pull down

Clones were generated using the Gateway cloning system (Thermo Fisher Scientific); the desired inserts were cloned into pDONR221 as pENTR clones and then into different destination vectors using the LR clonase recombination method (Thermo Fisher Scientific). We generated full-length ORF pENTR clones for AtFANCC, AtFANCE, and AtMHF2 from an inflorescence cDNA library of Arabidopsis. One additional ORF pENTR of AtFANCC was cloned without a stop codon for in-frame C-terminal fusion and both ORF pENTR clones of AtFANCC were used for GSrhino-tagged pulldown. In yeast two-hybrid assays, we used two destination vectors, pGADT7-GW as bait and pGBKT7-GW as prey. The ORF pENTR clones of AtFANCC, AtFANCE, and AtMHF2 were cloned into both destination vectors by LR reaction. The ORF of AtFANCF was cloned into the pGBKT7 and PACT2 AD convectional vector using the NCO1, Sal1, and NCO1, XHO1 restriction enzymes, respectively. All pENTR clones and final clones were verified thoroughly by Sanger sequencing to ensure mutation-free cloning and in-frame fusion. Plasmids of bait and prey clones were transformed into the haploid yeast strains AH109 and Y187, and then yeast two-hybrid assays were performed in a Gal4-based system from Clontech in a diploid strain by mating as previously described (Rossignol et al. 2007; Seguela-Arnaud et al. 2017).

Arabidopsis cell suspension cultures expressing N-terminal GSrhino-tagged FANCC and for C-terminal GSrhino-tagged FANCC were used for pull-down as previously described (Van Leene et al. 2019; Cromer et al. 2019). Co-purified proteins were identified using standard protocols utilizing on bead-digested sample evaluated on a Q Exactive mass spectrometer (Thermo Fisher Scientific) (Van Leene et al. 2015). After identification, the protein list was filtered for false-positives using a list of non-specific proteins, which was assembled as previously reported (Van Leene et al. 2015). Semi-quantitative analysis using the average normalized spectral abundance factors (NSAF) of the discovered proteins in the FANCC pull-downs was used to identify true interactors that may have been filtered out due to their classification in the list of nonspecific proteins. Chosen proteins were identified with at least two peptides in at least two experiments and showed high (at least 10-fold) and significant [log10(P value (t test)) enrichment relative to estimated average NSAF values from a large dataset of pull-downs with nonrelated bait proteins.

### FTL analysis

To measure recombination, we used fluorescent transgenic lines (FTL) (420) generated in a Col background. The used lines harbor seed coat expressing GFP (Chr 3:256,516-GFP) and dsRed (Chr 3:5,361,637-dsRed2) fluorescent protein markers in cis (Ziolkowski et al. 2015; Melamed-Bessudo et al. 2005). We quantified the fluorescence of the seeds using the Fiji image analysis software (Schindelin et al. 2012), which identifies seeds and quantifies fluorescence intensity for each seed in all pictures. The output was analyzed using a pipeline that was created to normalize the data, plot the frequency of objects with each fluorescent color, plot the fluorescence intensity, and quantify the number of seeds with only one fluorescent color, allowing selection of the number of recombinant seeds. For F2, recombination was measured using the formula below, as reported in (Ziolkowski et al. 2015).

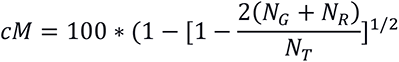

For male and female backcrosses, recombination was measured as

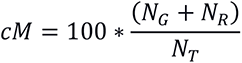

Where *N_G_* is the number of green-only fluorescent seeds, *N_R_* is the number of red-only fluorescent seeds and *N_T_* is the total number of seeds counted.

We generated a segregating population (F2) from which we selected plants heterozygous for the markers in cis with the desired mutants and wild-type control. For each genotype, we used at least three biological replicates (independent plants) with at least three technical replicates, each of them containing a minimum of 400 seeds. To measure CO frequency independently in males and females, reciprocal crosses were made with wild-type Columbia (0). Differences between genotypes were tested by Chi2 on the proportion of recombined seeds (Ng+Nr) among total seeds.

## Funding

This work was supported by core funding from the Max Planck Society (to R.M.), a Depart of Biotechnology Centre of Excellence grant (to I.S.), and the CEFIPRA project SMOKI (to R.M. and IS).

**Figure S1. Schematic representation of the *AtFANCC*, *AtFANCE* and *AtFANCF* genes.**

Gene orientations are indicated by horizontal arrows. A vertical red line indicates a point mutation, and red triangles indicate T-DNA insertions. The numbers on inverted triangles and vertical line denote corresponding alleles of *fancc*. Exons are indicated by a solid black box, while introns and untranslated regions (UTR) are represented by a line.

**Figure S2. Analysis of fertility of *zmm* suppreso*r* mutants**. Each dot indicates the fertility of an individual plant, measured as the number of seeds per fruit averaged on ten fruits. The mean for each genotype is represented by a red bar. The vertical lines separate experiments performed independently. Within each experiment, all plants were cultivated together in a population segregating for the mutations. *P* values are from one-way ANOVA followed by the Sidak test for multiple comparisons.

**Figure S3. Superimposition of the five models of the AtFANCC-AtFANCE-AtFANCF complex calculated by AlphaFold2.** (A) Representation showing the convergence of the five models of FANCC and FANCF superimposed with only one example of FANCE for the sake of clarity. (B) Same view as (A) with the five models of FANCE shown in different shades of pink color. In the top panel, the C-terminal orientation of FANCE is found to be poorly defined due to the flexible short linker connecting it to its N-terminal domain (lower panel) whose orientation is, in contrast, very well defined due to a large binding interface with FANCC. (C) 2D map of the predicted alignment error (PAE) calculated by AlphaFold2 which can be used as a proxy for the reliability of the structural model of every subunit and of the binding interfaces. As in panel B, the interface between the N-terminal domain of FANCE and FANCC is predicted to be accurately modeled, while its C-terminal domain is not predicted to bind as specifically and reliably. (D) Representation of the PAE map calculated for each of the five AlphaFold2 models.

**Figure S4. Multiple sequence alignment of FANCC orthologs in ten representative plant species.** The structural model of the complex is shown on top. The abbreviated species names and their NCBI indexes are provided in the headers. The regions of FANCC interacting with FANCF and FANCE are indicated by blue and pink straight lines around them, respectively. MSA is represented using JalView (Waterhouse et al. 2009).

**Figure S5. Circular representation of the phylogenetic tree of FANCC orthologs.** A phylogenetic tree comprising 72 plant species orthologs was calculated using PhyML (Dereeper et al. 2008) and represented with iTOL(Letunic and Bork 2021). A sampled multiple sequence alignment of nine FANCC orthologs in the species labelled at the end of the branches is available in Figure S4. Confidence supports for each clade in the tree are indicated by the size of the gray disks; the reliable branches are green, while the unreliable ones range from yellow to red.

**Figure S6**. FANCC interactions tested by yeast two-hybrid.

Proteins of interest were fused with Gal4 DNA binding domain (BD) as bait and with Gal4 activation domain as prey (AD), respectively, then expressed in yeast cells. For each combination, yeast cells were spotted on non-selective medium (-LW) as control and moderately selective media (-LWH). Growth on -LWH is interpreted as direct interaction between the two tested proteins.

**Figure S7**. Chromosome spreads of male meiocytes at metaphase I.

(A–B) *fance* (C–D) *fancf,*A and C are normal metaphase I with five bivalents while B and D have one pair of univalent and four bivalents. (Scale bar, 10 µm).

**Figure S8. Analysis of fertility of *fanc* mutants combined with *mus81***. Each dot indicates the fertility of an individual plant, measured as the number of seeds per fruit averaged on ten fruits. The mean for each genotype is represented by a red bar. The vertical lines separate independent experiments. All plants were cultivated in parallel in each experiment, and the wild-type controls were mutant siblings except for the *mus81 fance* combination because the two genes are linked. In this latter case, *mus81* segregating and *fance* segregating population mutants were used as controls. *P* values are from one-way ANOVA followed by the Sidak test for multiple comparisons.

**Figure S9. Combining *fanc* and *mus81* mutations instigates chromosome fragmentation at meiosis.** (A–K) Chromosome spreads of male meiocytes (Scale bar, 10 µm). Arrow heads indicate chromosome fragments.

Table S1. Mutations identified in the *zmm* suppressor screen.

Table S2. FTL seed data used to measure genetic recombination.

Table S3. Primers used in this study.

Table S4. MS data

## Acknowledgements

We thank Virginie Portemer for her help in *fancc-1* mapping. We thank Piotr A. Ziolkowski for kindly providing the 420 FTL line. We thank the VIB proteomics core facility for performing the Q Exactive analysis of the pull down samplesand Neysan Donnelly for proofreading the manuscript.

## Notes

### Competing Interest Statement

The authors have declared no competing interest.

