## supplemental figures for "The FANCC-FANCE-FANCF complex is evolutionarily conserved and regulates meiotic recombination"

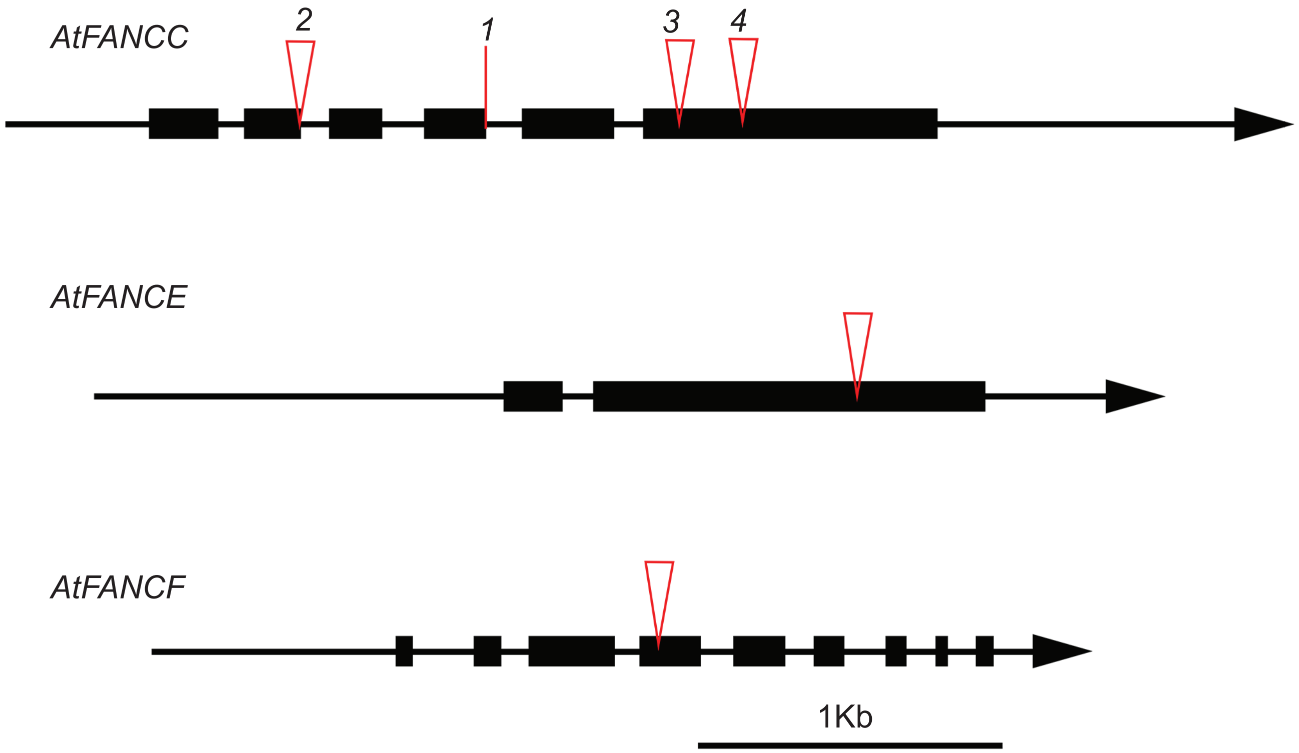


**Figure S1. Schematic representation of the *AtFANCC, AtFANCE* and *AtFANCF* genes.**
Gene orientations are indicated by horizontal arrows. A vertical red line indicates a point mutation, and red triangles indicate T-DNA insertions. The numbers on inverted triangles and vertical line denote corresponding alleles of fancc. Exons are indicated by a solid black box, while introns and untranslated regions (UTR) are represented by a line.

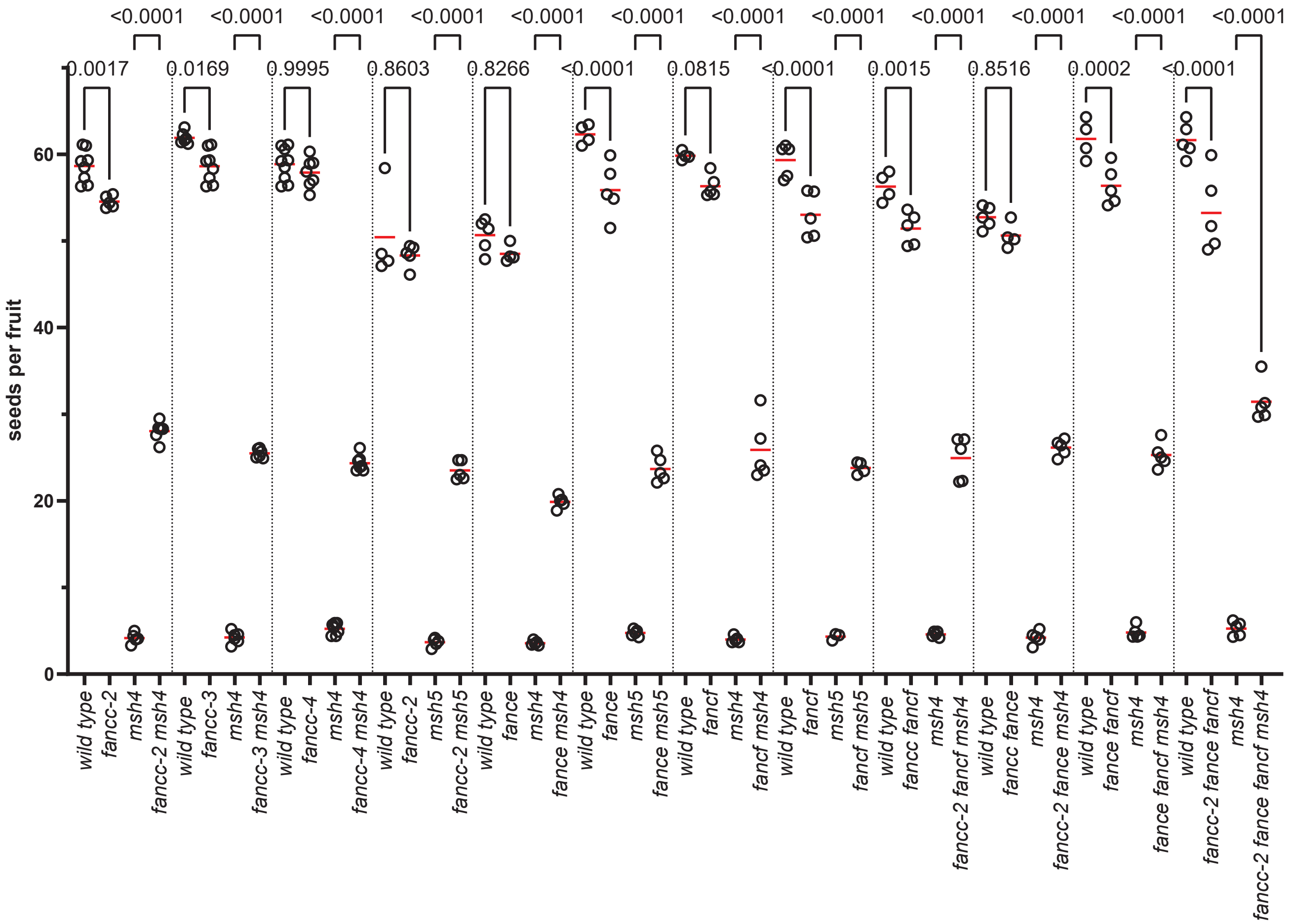


**Figure S2. Analysis of fertility of zmm suppresor mutants.** Each dot indicates the fertility of an individual plant, measured as the number of seeds per fruit averaged on ten fruits. The mean for each genotype is represented by a red bar. The vertical lines separate experiments performed independently. Within each experiment, all plants were cultivated together in a population segregating for the mutations. P values are from one-way ANOVA followed by the Sidak test for multiple comparisons.

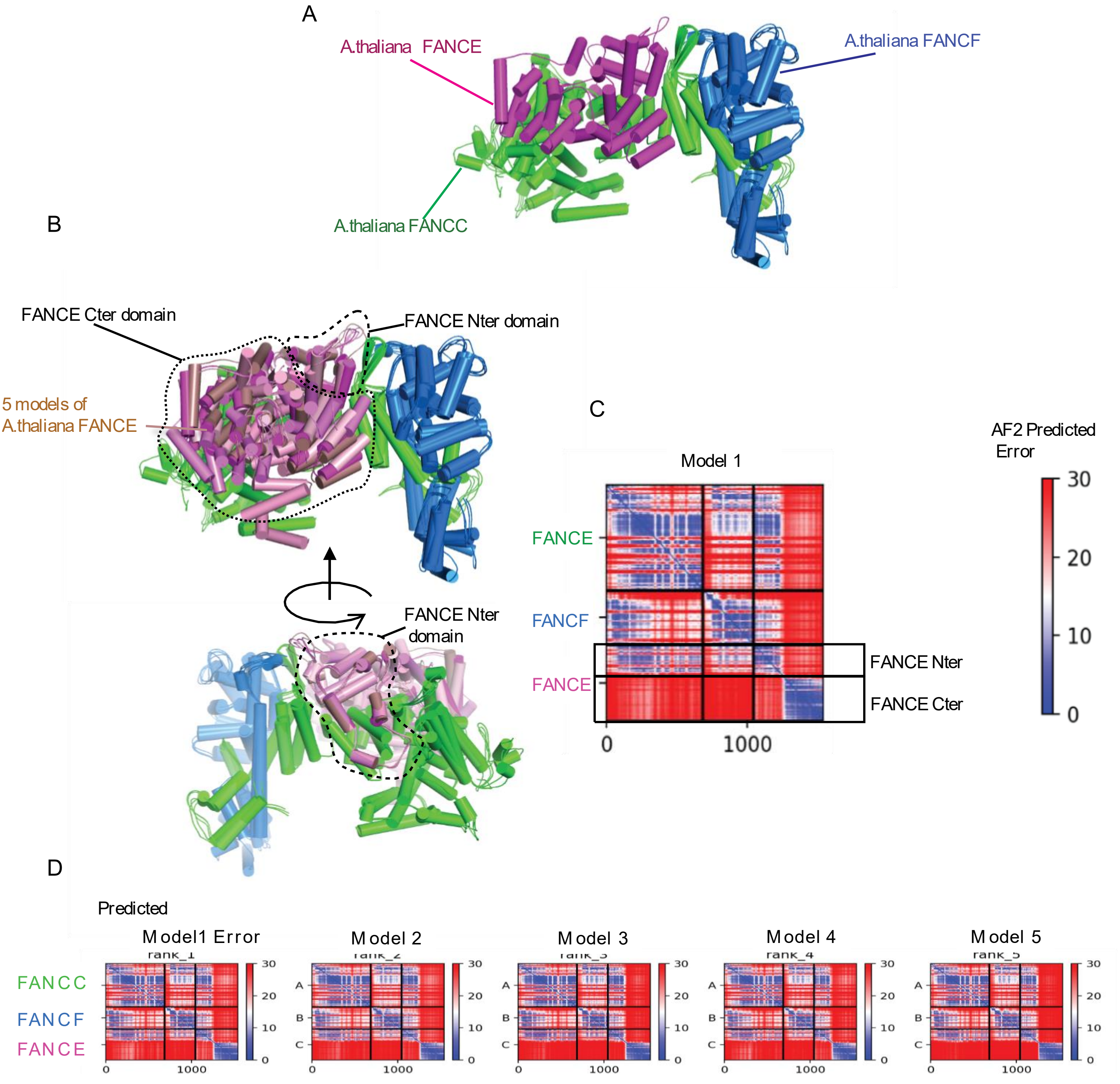


**Figure S3. Superimposition of the five models of the AtFANCC-AtFANCE-AtFANCF complex calculated by AlphaFold2.** (A) Representation showing the convergence of the five models of FANCC and FANCF superimposed with only one example of FANCE for the sake of clarity. (B) Same view as (A) with the five models of FANCE shown in different shades of pink color. In the top panel, the C-terminal orientation of FANCE is found to be poorly defined due to the flexible short linker connecting it to its N-terminal domain (lower panel) whose orientation is, in contrast, very well defined due to a large binding interface with FANCC. (C) 2D map of the predicted alignment error (PAE) calculated by AlphaFold2 which can be used as a proxy for the reliability of the structural model of every subunit and of the binding interfaces. As in panel B, the interface between the N-terminal domain of FANCE and FANCC is predicted to be accurately modeled, while its C-terminal domain is not predicted to bind as specifically and reliably. (D) Representation of the PAE map calculated for each of the five AlphaFold2 models.

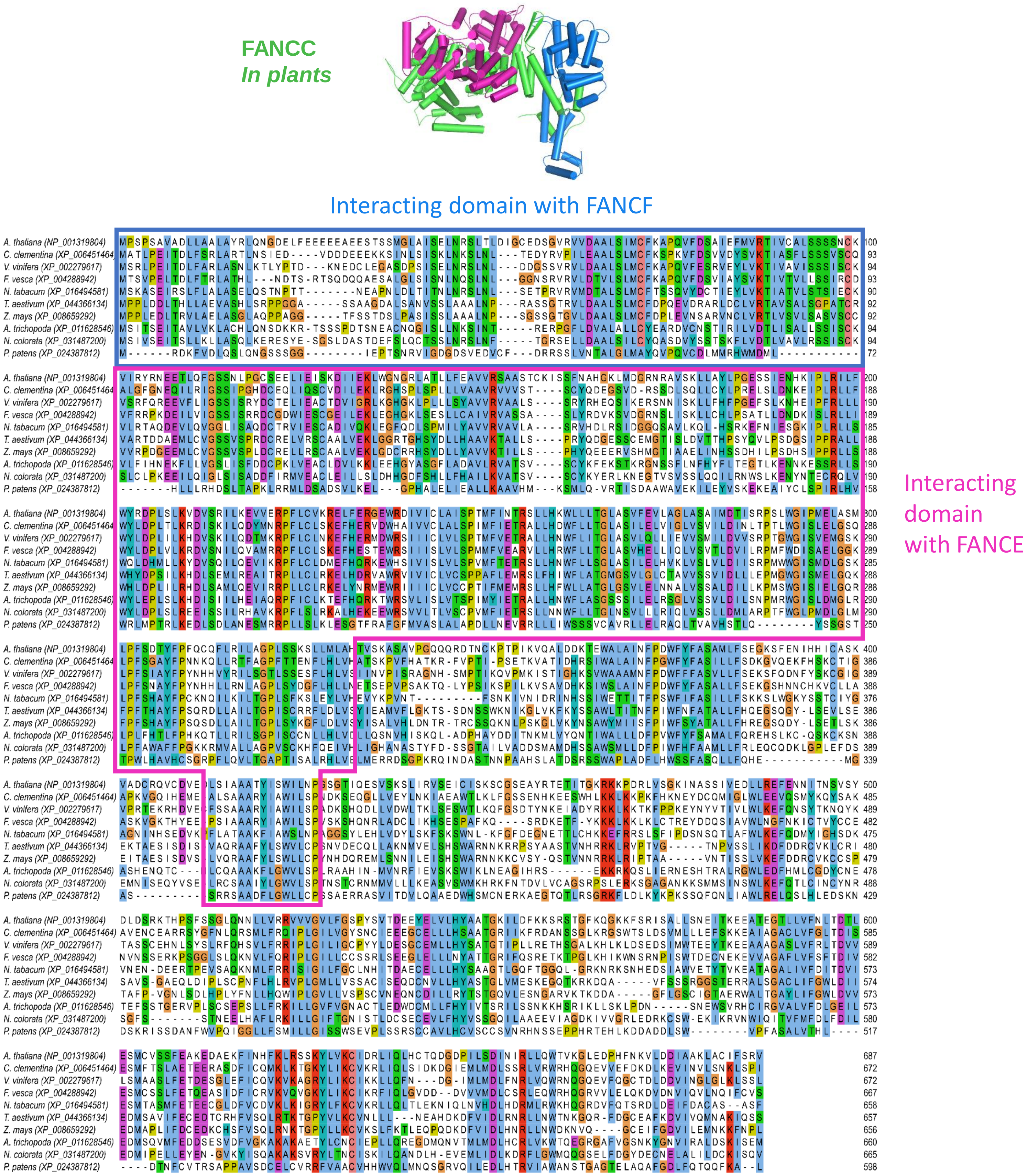


**Figure S4. Multiple sequence alignment of FANCC orthologs in ten representative plant species.** The structural model of the complex is shown on top. The abbreviated species names and their NCBI indexes are provided in the headers. The regions of FANCC interacting with FANCF and FANCE are indicated by blue and pink straight lines around them, respectively. MSA is represented using JalView (Waterhouse et al. 2009).

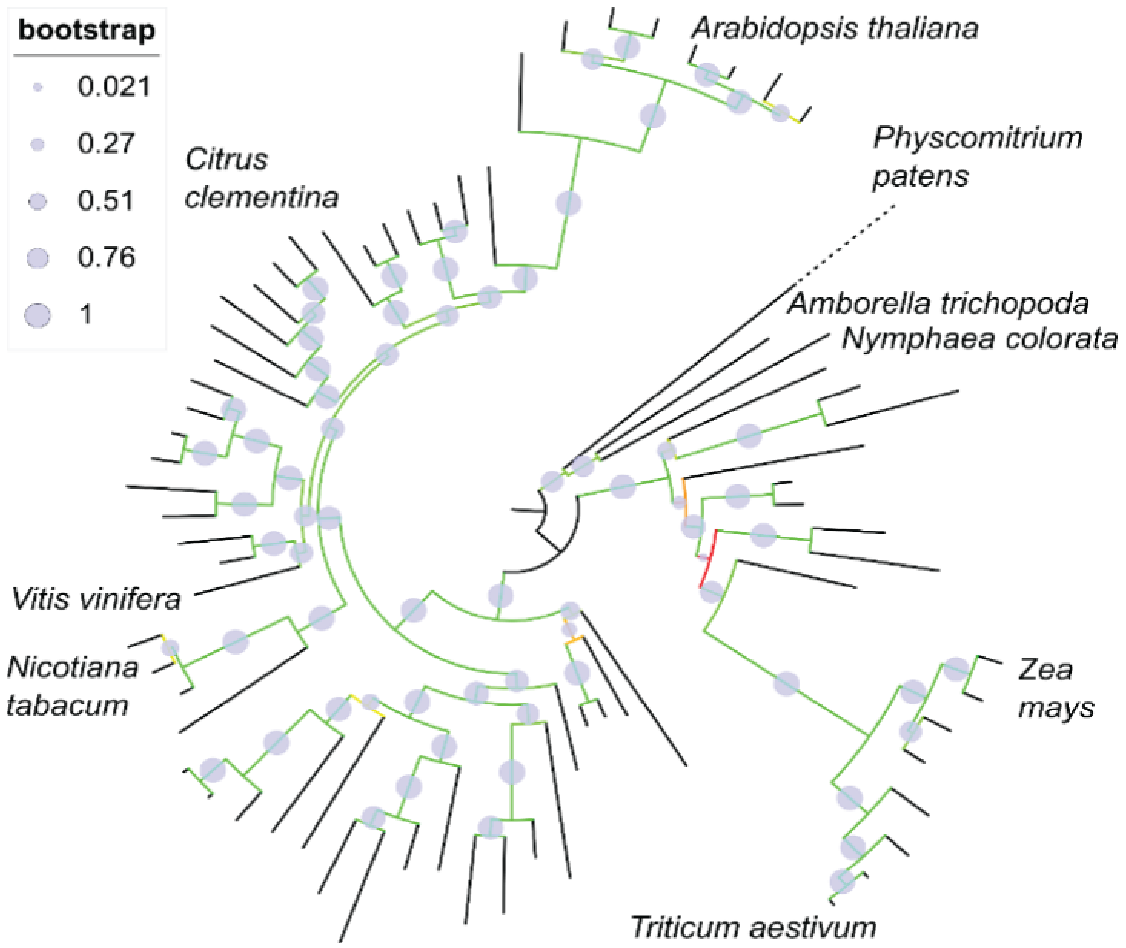


**Figure S5. Circular representation of the phylogenetic tree of FANCC orthologs.** A phylogenetic tree comprising 72 plant species orthologs was calculated using PhyML (Dereeper et al. 2008) and represented with iTOL (Letunic and Bork 2021). A sampled multiple sequence alignment of nine FANCC orthologs in the species labelled at the end of the branches is available in Figure S4. Confidence supports for each clade in the tree are indicated by the size of the gray disks; the reliable branches are green, while the unreliable ones range from yellow to red.

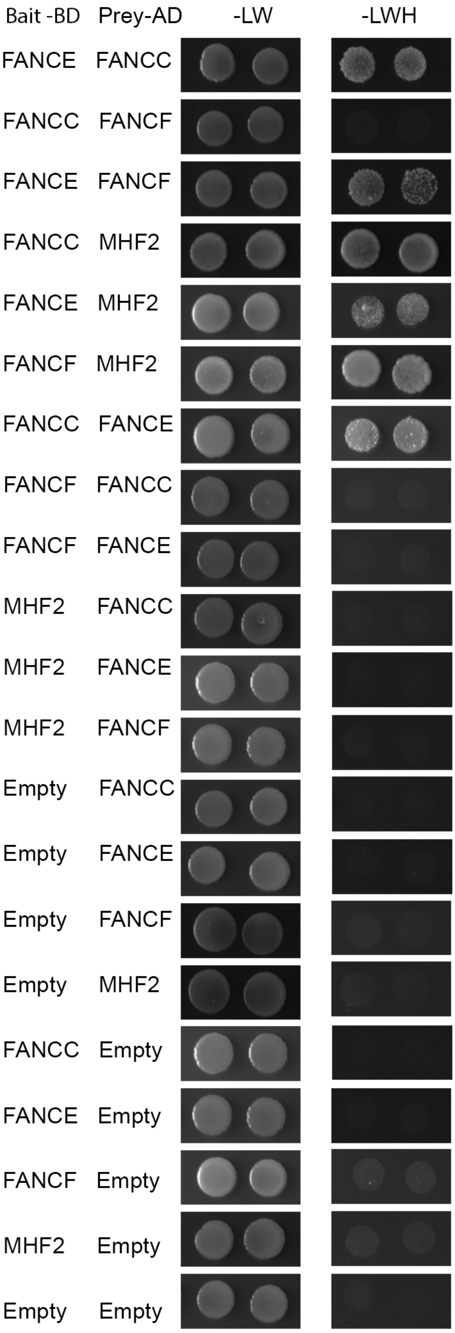


**Figure S6. FANCC interactions tested by yeast two-hybrid.**

Proteins of interest were fused with Gal4 DNA binding domain (BD) as bait and with Gal4 activation domain as prey (AD), respectively, then expressed in yeast cells. For each combination, yeast cells were spotted on non-selective medium (-LW) as control and moderately selective media (-LWH). Growth on -LWH is interpreted as direct interaction between the two tested proteins.

.

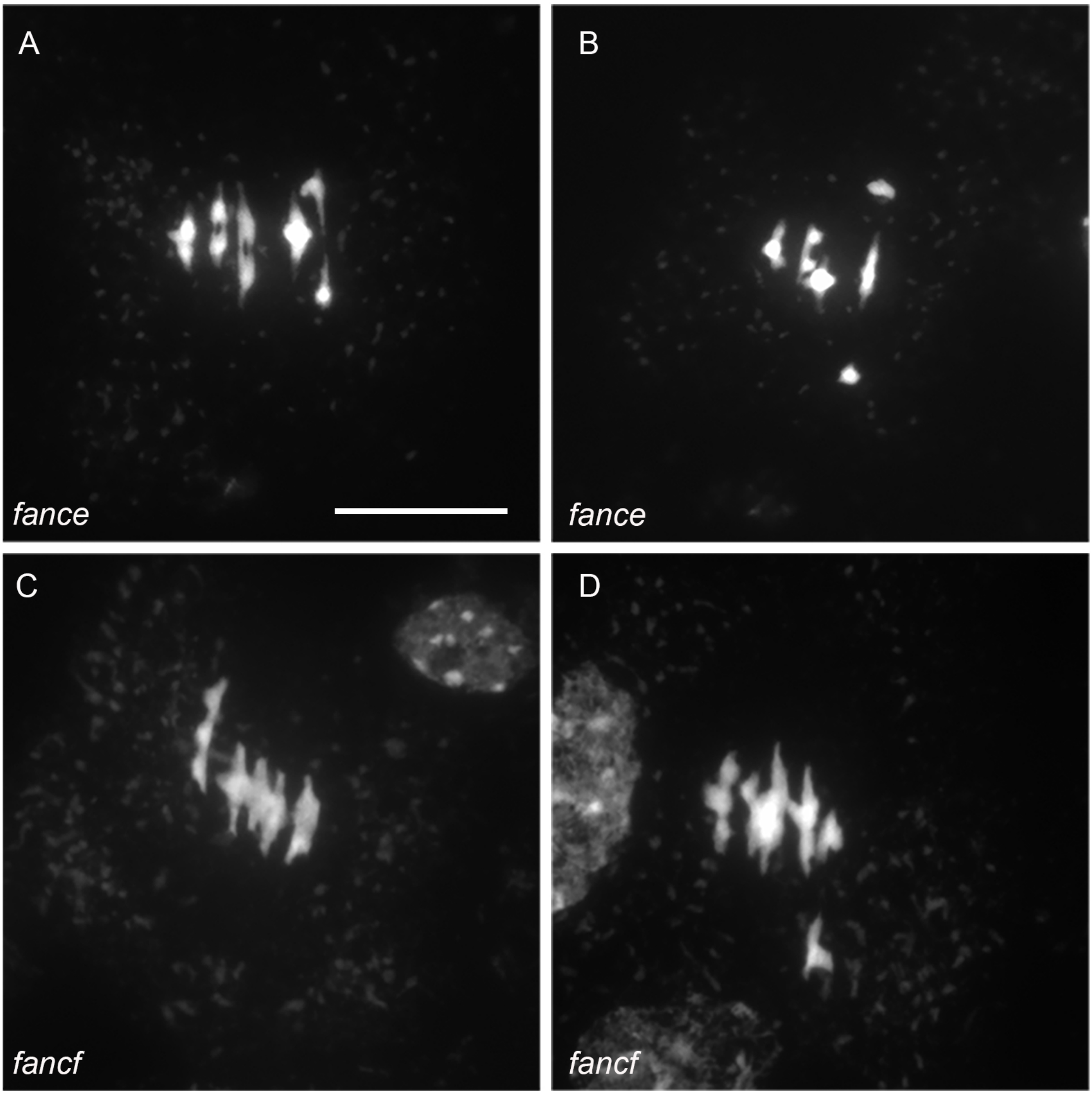


**Figure S7. Chromosome spreads of male meiocytes at metaphase I.** (A–B) fance (C–D) fancf,A and C are normal metaphase I with five bivalents while B and D have one pair of univalent and four bivalents. (Scale bar, 10 μm).

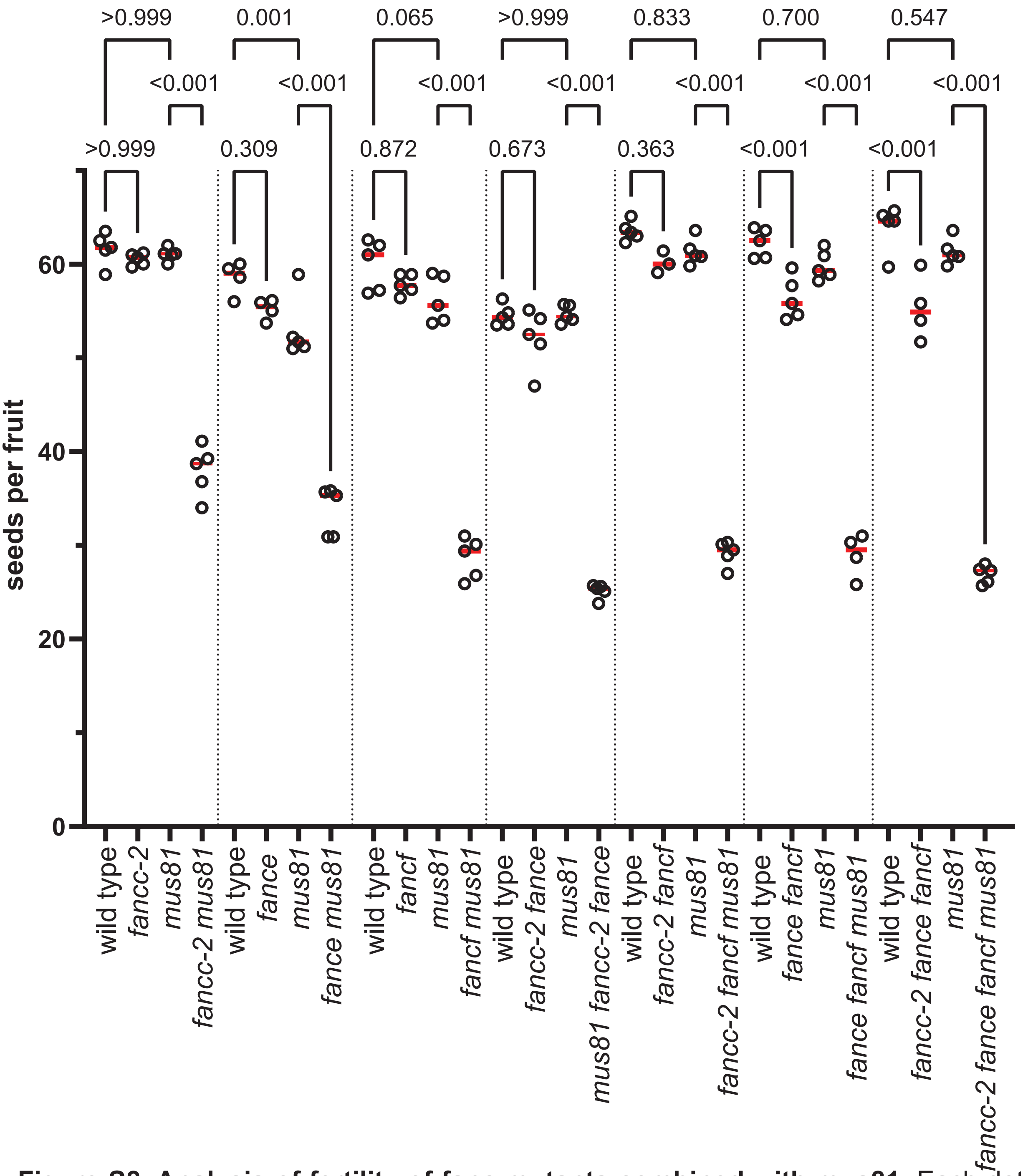


**Figure S8. Analysis of fertility of fanc mutants combined with mus81.** Each dot indicates the fertility of an individual plant, measured as the number of seeds per fruit averaged on ten fruits. The mean for each genotype is represented by a red bar. The vertical lines separate independent experiments. All plants were cultivated in parallel in each experiment, and the wild-type controls were mutant siblings except for the mus81 fance combination because the two genes are linked. In this latter case, mus81 segregating and fance segregating population mutants were used as controls. P values are from one-way ANOVA followed by the Sidak test for multiple comparisons.

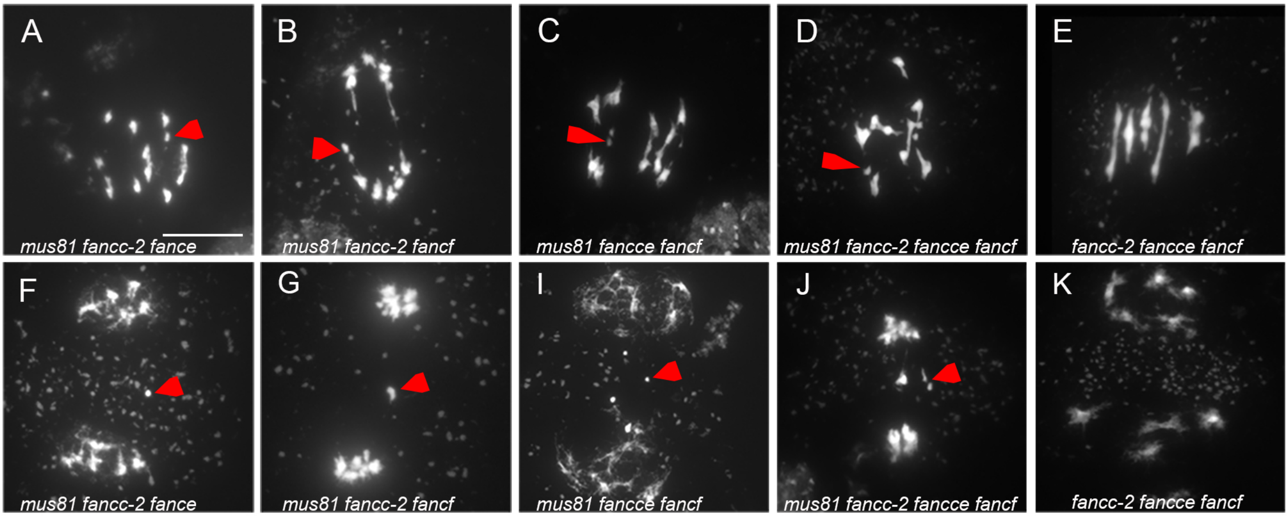


**Figure S9. Combining fanc and mus81 mutations instigates chromosome fragmentation at meiosis.** (A–K) Chromosome spreads of male meiocytes (Scale bar, 10 μm). Arrow heads indicate chromosome fragments.
